# Conifers concentrate large numbers of NLR immune receptor genes on one chromosome

**DOI:** 10.1101/2023.10.20.563305

**Authors:** Yannick Woudstra, Hayley Tumas, Cyril van Ghelder, Tin Hang Hung, Joana J. Ilska, Sebastien Girardi, Stuart A’Hara, Paul McLean, Joan Cottrell, Joerg Bohlmann, Jean Bousquet, Inanc Birol, John A. Woolliams, John J. MacKay

## Abstract

Nucleotide-binding domain and Leucine-rich Repeat (NLR) immune receptor genes form a major line of defence in plants, acting in both pathogen recognition and resistance machinery activation. NLRs are reported to form large gene clusters in limber pine (*Pinus flexilis*) but it is unknown how widespread this genomic architecture may be among the extant species of conifers (Pinophyta). We used comparative genomic analyses to assess patterns in the abundance, diversity and genomic distribution of NLR genes. Chromosome-level whole genome assemblies and high-density linkage maps in the Pinaceae, Cupressaceae, Taxaceae and other gymnosperms were scanned for NLR genes using existing and customised pipelines. Discovered genes were mapped across chromosomes and linkage groups, and analysed phylogenetically for evolutionary history. Conifer genomes are characterised by dense clusters of NLR genes, highly localised on one chromosome. These clusters are rich in TNL-encoding genes, which seem to have formed through multiple tandem duplication events. In contrast to angiosperms and non-coniferous gymnosperms, genomic clustering of NLR genes is ubiquitous in conifers. NLR-dense genomic regions are likely to influence a large part of the plant’s resistance, informing our understanding of adaptation to biotic stress and the development of genetic resources through breeding.

**Plain language summary:** NLR immune receptor genes are important in pest, disease and drought resistance of plants. In the giga-genomes of conifers, they concentrate on very small chromosomal regions. These regions act as important reservoirs for NLR diversity and can be used in breeding to improve the resilience of conifer trees.

## Introduction

Disease resistance is one of the key areas of study in plant genetics and evolution with implications for conservation and ecosystem health, as well as breeding. Decades of research have improved our understanding of the identity and interplay of major gene families involved in disease resistance with functions in detection and defence (Ngou, Ding, and Jones 2022). One of the first events in plant defence mechanisms is pathogen recognition, in which the Nucleotide-binding domain and Leucine-Rich Repeat (NBS-LRR or NLR) immune receptor gene family plays a central role (Duxbury, Wu, and Ding 2021). The products of NLR genes occur intracellularly and can bind directly to specific pathogenic effectors (pathogen-encoded proteins) or detect modifications of plant proteins induced by such effectors, thus activating a cascade of defence mechanisms upon perception (Ngou, Ding, and Jones 2022). NLRs provide a typical example of an evolutionary arms race in which pathogenic effectors evolve to evade detection by host NLRs which, in turn, evolve to recognise the new variants (e.g., the *Capsicum chinense* Jacq. NLR *Tsw* versus the pathogen ‘Tomato spotted wilt virus’ (Chen et al. 2023)). Unsurprisingly therefore, NLRs are diverse and abundant in many plant species with several hundred different NLR genes found in a range of land plant lineages (Barragan and Weigel 2021). While NLRs have been studied extensively in angiosperms (i.e, Cucurbitaceae (Lin et al. 2013), Rosaceae (Jia et al. 2015) and Solanaceae (Seo et al. 2016); Angiosperm NLR Atlas (Y. Liu et al. 2021)), studies in conifers are rare (Van Ghelder et al. 2019; J.-J. Liu et al. 2019; Ence et al. 2022).

NLRs exhibit a conserved tripartite structure consisting of 1) a non-conserved N-terminal domain; 2) a conserved central nucleotide-binding domain (NB-ARC, defined as “a nucleotide-binding adaptor shared by APAF-1, certain *R* gene products and CED-4” (van der Biezen and Jones 1998)); and 3) a C-terminal leucine-rich repeat (LRR) domain which can vary in length. These resistance genes seem to have originated before the rise of land plants (Shao et al. 2019) and have since diversified into three main classes based on the character of the N-terminal domain: CNL, RNL and TNL. CNLs (N-terminal ‘coiled-coil’ domain) and RNLs (N-terminal ‘resistance to powdery mildew 8 (RPW8) domain’) are closely related classes, whereas TNLs (N-terminal ‘Toll interleukin-1 receptor’ domain) form a distinct class. All three classes are unusually abundant and diverse in conifers, with an RNL diversity that is distinctly higher than in any other group of land plants (Van Ghelder et al. 2019). Furthermore, a genomic distribution analysis in limber pine (*Pinus flexilis* E.James) revealed an unbalanced intragenomic and intrachromosomal distribution of NLRs (J.-J. Liu et al. 2019). In this species, one chromosome contained a dense cluster of NLRs comprising mainly TNLs, indicating a high rate of tandem duplications. Besides disease resistance, NLRs have been shown to be responsive to drought stress when investigated in the conifer white spruce (*Picea glauca* (Moench) Voss) (Van Ghelder et al. 2019). Controlling both disease and drought resistance, NLR-rich genomic regions will be particularly interesting candidates for genomic breeding purposes and genetic resource management in conifers, especially under climate change and the spread of tree pests and diseases.

Conifers are found in diverse ecosystems and several species are used in productive forestry across the globe, some of which involves breeding programs (Mullin et al. 2011). Breeding in conifers traditionally relies on pedigree analysis and phenotypic evaluations such as growth rates, wood yield and properties, and susceptibility to biotic and abiotic threats (White, Adams, and Neale 2007). Emerging and intensifying threats to the health of conifers in natural populations and managed forests involve a range of biotic stressors including oomycetes, fungi, herbivorous insects, and nematodes (Mitton and Ferrenberg 2012; Jakoby, Lischke, and Wermelinger 2019; Brar et al. 2018; Mota et al. 1999). In response to these challenges, molecular tools and genomic resources are being developed to support both fundamental research and diverse applications such as an acceleration of breeding outputs (Neale and Kremer 2011; Stocks et al. 2019; Bousquet et al. 2021). For example, investigations have linked genetic resistance to fusiform rusts in *Pinus taeda* L. to TNL-encoding sequences and to genomic clustering of resistance genes (Wilcox et al. 1996; Quesada et al. 2014; Ence et al. 2022). Genomic selection methods may have the potential to be developed to enhance resistance, such as rust resistance in *Pinus taeda* L. (Ence et al. 2022) and insect resistance in Norway spruce (*Picea abies* (L.) H.Karst.) (Lenz et al. 2020). However, the success of this relies on a more complete understanding of the genomic architecture of conifer NLRs (Ence et al. 2022).

The large size (often ≥ 10 Gbp) and relative complexity of conifer nuclear genomes has challenged whole genome sequence assembly (e.g., Nystedt et al. 2013; Birol et al. 2013; Zimin et al. 2014) but new methods have greatly improved the contiguity of genome assemblies as seen in *Sequoiadendron giganteum* (Lindl.) J.Buchholz (Scott et al. 2020), *Taxus chinensis* (Pilg.) Rehder (Xiong et al. 2021), and *Pinus tabuliformis* Carrière (Niu et al. 2022). In some species that lack such assemblies, high-density genetic maps are available to probe genome architecture (J.-J. Liu et al. 2019; Bernhardsson et al. 2019; Gagalova et al. 2022). Together, these assembled genome sequences and genetic maps, along with diverse transcriptomes (e.g., Van Ghelder et al. 2019) open the doors to more comprehensive analyses of resistance genes and their genomic architecture in conifer trees. Considering the relatively high level of genome conservation across conifers, we may predict that genomic clustering as observed in *P. flexilis* (J.-J. Liu et al. 2019) will occur across conifer taxa, that is to the extent they result from shared ancestral evolutionary events.

Considering the potential benefits of genomic breeding with NLR dense genome segments for conservation and industry, we investigated NLR gene clustering patterns across conifers. We leveraged recently published diploid high-density linkage maps and chromosome-level whole genome assemblies for genomic mapping of NLR genes. Results for conifers were contrasted with non-conifer gymnosperms from the Ginkgoales and Cycadales (Figure 1). To elucidate the evolutionary trajectories towards the observed clustering patterns, NLR genes were analysed in a phylogenetic framework. Our results indicate consistently uneven genomic distribution patterns of NLR genes across all conifers, with large and heavily concentrated reservoirs of NLR genes located on specific chromosomes. This knowledge on resistance genes will be informative for both breeding and conservation of these economically and ecologically important trees.

**Figure 1:**
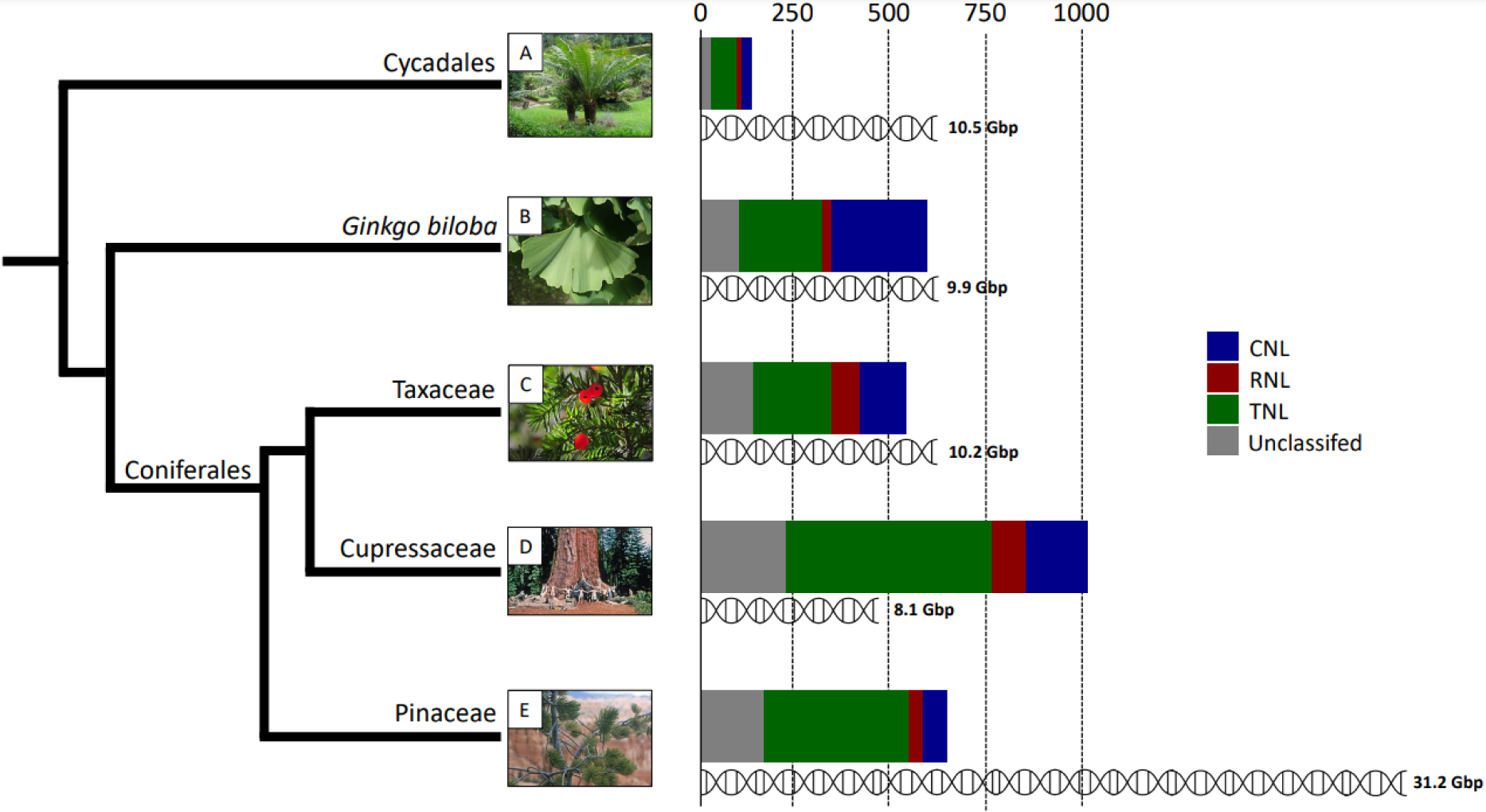
Overview of main gymnosperm clades, with total number of NLR genes and their corresponding categories indicated as bar charts, as found in this study (Table 1). Cladogram is based on the phylogeny presented in Leslie et al. (2018). Genome sizes are indicated beneath each bar chart and are based on the chromosome-level assemblies used in this study (Materials & methods section 1 for details) or, in the case of Pinaceae (*Pinus flexilis*), obtained from the Kew Plant DNA C-values database (release 7.1, Pellicer and Leitch 2020). Gymnosperms invariably have large genomes (∼10 Gbp), but display large variations in NLR gene numbers. Despite the three-fold increase in genome size observed in Pinaceae, the number of discovered NLR genes remained within the average range of conifers. Pictures were obtained from the Wikimedia Commons repository (https://commons.wikimedia.org) and correspond to the broader taxonomic clades: A – *Cycas rumphii* Miq., andy_king50; B – *Ginkgo biloba* L., Susanna Giaccai; C – *Taxus baccata* L., Mykola Swarnyk; D – *Sequoiadendron giganteum*, W. Bulach; E – *Pinus flexilis*, Greg Woodhouse.

**Table 1:** Number of NLR genes discovered in each taxon of this study, separated by class. As some NLR genes were located on unassigned scaffolds in *C. panzhihuaensis*, *G. biloba* and *T. chinensis*, the numbers of NLR genes mapped onto chromosomes are recorded separately in brackets alongside the total number of NLR genes discovered for these species. Some RNL comprised genes only in the RPW8 domain, and these are recorded separately in brackets for each taxon. \**P. flexilis* values are based on the results from J.-J. Liu et al (2019). **Values for *Picea* taxa are based on NLR annotation of high-density linkage maps or incomplete genome assemblies that were built using high-density linkage maps (more details are provided in Materials & methods sections section 1 & 2). ^Listed only for the chromosome with the highest NLR-density. The method for calculating the ratio of observed /predicted NLR clustering on a single chromosome is explained in Materials & methods section 1.

| *Species* | *Total #NLR*<br>*(on chr.)* | *#CNL* | *#RNL*<br>*(RPW8 only)* | *#TN*<br>*L* | *Uncl*<br>*.* | *Densest chr.*<br>*(% of total)* | *#Observed*<br>*/#Predicted^* | *p-value*<br>*(distrib.)* |
| --- | --- | --- | --- | --- | --- | --- | --- | --- |
| *Cycas panzhihuaensis* | 136 (127) | 29 | 13 (8) | 65 | 29 | 24 (19%) | 2.02 | 0.02958 |
| *Ginkgo Biloba* | 585 (570) | 249 | 25 (5) | 213 | 98 | 83 (14%) | 2.04 | 6.70e-08 |
| ***Taxaceae*** |  |  |  |  |  |  |  |  |
| *Taxus chinensis* | 533 (496) | 124 | 73 (29) | 201 | 135 | 146 (29%) | 4.30 | < 2.2e-16 |
| ***Cupressaceae*** |  |  |  |  |  |  |  |  |
| *Sequoiadendron giganteum* | 1,002 | 159 | 89 (17) | 533 | 221 | 312 (31%) | 2.47 | < 2.2e-16 |
| ***Pinaceae*** |  |  |  |  |  |  |  |  |
| *Pinus flexilis* * | 639 | 64 | 36 (-) | 376 | 163 | 266 (42%) | 3.55 | < 2.2e-16 |
| *Picea abies* ** | 173 | 37 | 30 (5) | 85 | 21 | 68 (39%) | 3.37 | 4.78e-05 |
| *Picea glauca* ** | 273 | 74 | 43 (21) | 105 | 51 | 89 (33%) | 3.26 | 1.07e-07 |
| *Picea sitchensis* ** | 245 | 73 | 50 (26) | 81 | 41 | 57 (23%) | 2.61 | 0.003825 |

## Results

NLR immune receptor gene diversity and abundance were annotated in the genomes of six conifer species (members of the Pinaceae, Cupressaceae and Taxaceae) and two other gymnosperms (*Ginkgo biloba* and a member of the Cycadales), with varying levels of contiguity. We deployed an automated NLR annotation pipeline and additional manual BLAST procedures on recently published high-density linkage maps and chromosome-level whole genome assemblies to physically map the genomic distribution of NLR genes in gymnosperms. We discovered consistent patterns of genomic clustering of NLR genes among conifers, but not in other gymnosperms. In each of the analysed conifers, a particularly dense cluster of NLR genes occurred on a single chromosome which contained between 18% and 34% of the total number of NLR genes within only a short segment of a few Mbp (or cM in the case of linkage maps).

We restricted our analysis to diploid genomes to avoid potential issues of false discovery for duplicated genes, due to incomplete phasing of the haplotypes in polyploid genome assemblies. We therefore omitted the genome assembly of the hexaploid *Sequoia sempervirens* Endl. (Neale et al. 2022) from our analysis.

Our scan of genome assemblies and high-density linkage maps for conifers and other gymnosperms did include the recently published mega-genome assembly (25.4 Gbp) of the diploid *Pinus tabuliformis* (Niu et al. 2022), where some chromosomes surpass 2 Gbp in length. Unfortunately, the NLR Annotator pipeline (Steuernagel et al. 2020) is currently insufficiently programmed for such large contigs, limiting the output of our NLR analysis on this genome assembly. Furthermore, our preliminary results indicated extremely high NLR numbers (>4000) in this assembly, which we hypothesise to be an overestimation, considering the large disjunction with other conifers (1,002 in *S. giganteum*) and members of the *Pinus* genus (639 in *P. flexilis*). We therefore omitted *P. tabuliformis* from further analysis.

### 1. Gymnosperm NLR abundance and diversity

The number of NLR genes discovered in conifer genomic datasets varied nearly two-fold (Figure 1), between 533 (*Taxus chinensis*) and 1002 (*Sequoiadendron giganteum*). Of the non-conifers, *Ginkgo biloba* contained 585 NLR genes, which is similar to the average found in conifers (Table 1). In contrast, there were only 136 NLR genes found in the Cycad genome. Of the conifers, the *Sequoiadendron giganteum* genome contained the highest number of NLR genes (1,002) followed by (*Pinus flexilis*) which had considerably less (639). In all conifers, the TNL class was the most abundant, contributing between 33% (81/245, *Picea sitchensis*) to 59% (376/639, *Pinus flexilis*) to the total number of NLR genes (Table 1). In contrast, the CNL class was most abundant in *Ginkgo biloba*, in which they contributed 43% to the total number of NLR genes. RNLs were more abundant in conifers than in other gymnosperms. This class was proportionally most abundant in *Picea* species (16-20%), with half of the RNLs in *P. glauca* and *P. sitchensis* consisted only of RPW8 domains. The number of unclassified NLR genes (missing or ambiguous C-terminal domain) grew proportionally with the total number of NLR genes, representing 16-25% of the genes in complete genomic datasets.

### 2. Chromosomal and Intrachromosomal NLR distributions

In each of the conifer species we analysed, one chromosome displayed a disproportionately high NLR content (Table 1 and Figure 2A), a phenomenon that was absent in non-coniferous gymnosperms. In conifers, the chromosome which displayed the highest clustering of NLR genes contained between 29% (*T. chinensis*) and 42% (*P. flexilis*) of the total number of NLR genes found in the respective genome (Table 1). Even when there was incomplete genome coverage, the results for *Picea* species were within this range, the only outlier being *P. sitchensis* at 23%. Large numbers of NLR genes form dense clusters on these NLR-rich chromosomes or linkage groups (Figure 2B-E). These clusters contain high proportions of the total amount of NLR genes found in the genomic dataset, with up to 34% (*Pinus flexilis*, Figure 2C) of NLR genes concentrated in the space of 21% (44 cM) of a chromosome (linkage group).

**Figure 2:**
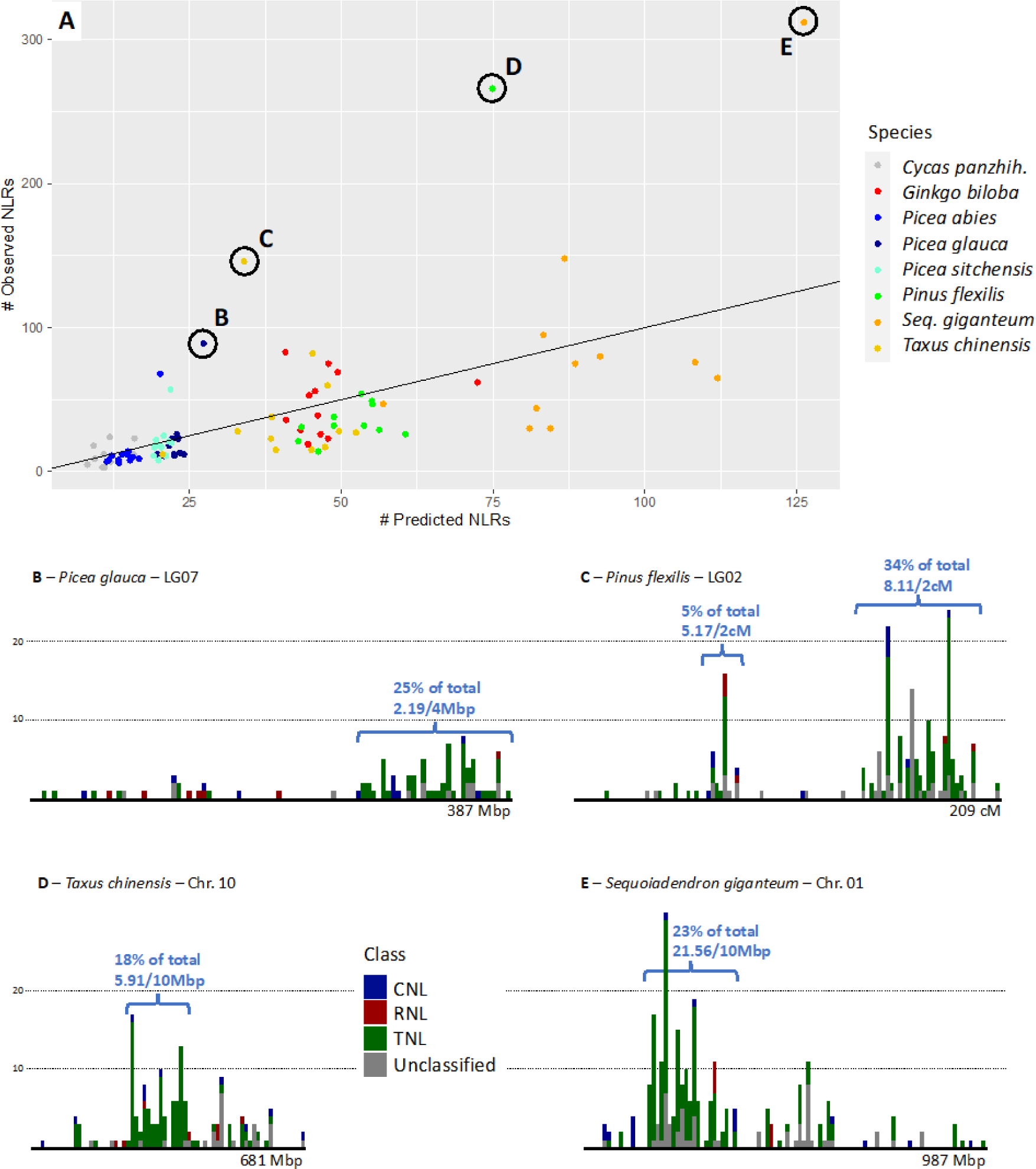
Chromosomal distribution of NLR genes across gymnosperms. A – Scatter plot indicating the observed number of NLR genes versus the expected number of NLR genes (calculated based on the length of the chromosome and the total number of NLR genes discovered in the genome, Materials & methods section 1) for each chromosome in each taxon analysed in this study. Black line follows the function y=x, indicating a perfectly homogeneous distribution of NLR genes over the chromosomes. Deviations from this line therefore indicate a non-homogeneous distribution. Highly deviant chromosomes of four taxa are highlighted and have their intrachromosomal NLR distribution displayed in histograms (B-E). Bin width equals ±1% of the length of the largest chromosome in the genome of the respective taxon (Materials & methods section 1). Colours indicate NLR class as determined with NLR Annotator (Steuernagel et al. 2020) and manual BLASTs (Materials & methods section 2). Ultra-dense NLR clusters are indicated for each taxon.

Although the distribution of NLR genes was non-random in all gymnosperms analysed, it was considerably less clustered in *C. panzhihuaensis* and *G. biloba* compared to the full genomic datasets of the conifers we analysed (Table 1). In the non-conifer species, only one small cluster of 13 genes (10% of total) was found on chromosome #10 in *C. panzhihuaensis* (Figure 3) and a small cluster of 45 genes (7.9% of total) occurred on chromosome #3 in *G. biloba,* both proportionally smaller than the clusters in conifer species (on average 25% of total). Non-uniform distribution of NLR genes in conifers did not only occur between but also within chromosomes (Figure 3) with large regions of NLR-rich chromosomes being devoid of NLR genes, particularly in *Taxus chinensis*.

**Figure 3:**
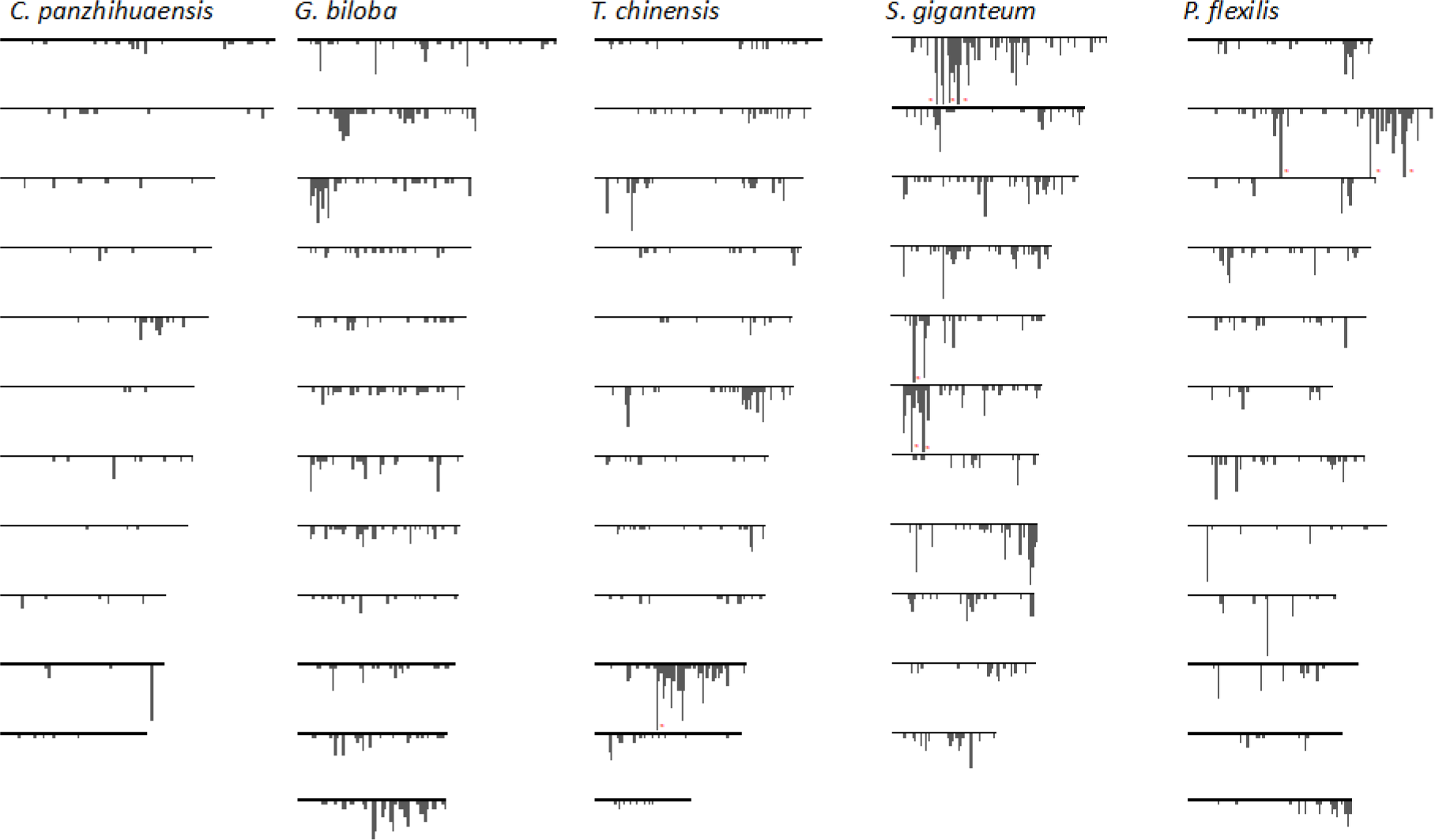
Histogram plots of NLR genes on each chromosome within the genome of five gymnosperm lineages. Chromosomes are ordered based on the ordering in the respective assembly and do not reflect synteny. Red **\*** indicates that a particularly dense bin has been cropped manually in order to fit all the histograms into a single image.

### 3. Intragenomic diversification and evolution of NLR genes

Phylogenetic relationships between intragenomic NLR genes followed expected class distinctions in all conifers and *Cycas panzhihuaensis* but not in *Ginkgo biloba* (Figure 4). Although conifer TNLs show a strong overall monophyletic correlation, *Taxus chinensis* (Fig. 4D) is the only gymnosperm where all TNLs share one most recent common ancestor (MRCA). *Sequoiadendron giganteum* (Fig. 4C) even has a small monophyletic clade of TNLs nested within the larger CNL/RNL clade. In all non-cycad gymnosperms, there is at least one monophyletic RNL clade with a different MRCA than the main CNL clade. All conifer CNL clades contained a few derived RNL sequences restricted to one phylogenetic subclade of the CNL clade. Both the *T. chinensis* and *S. giganteum* genomes contain a smaller monophyletic clade with a unique MRCA that contains almost exclusively unclassified NLR genes.

**Figure 4:**
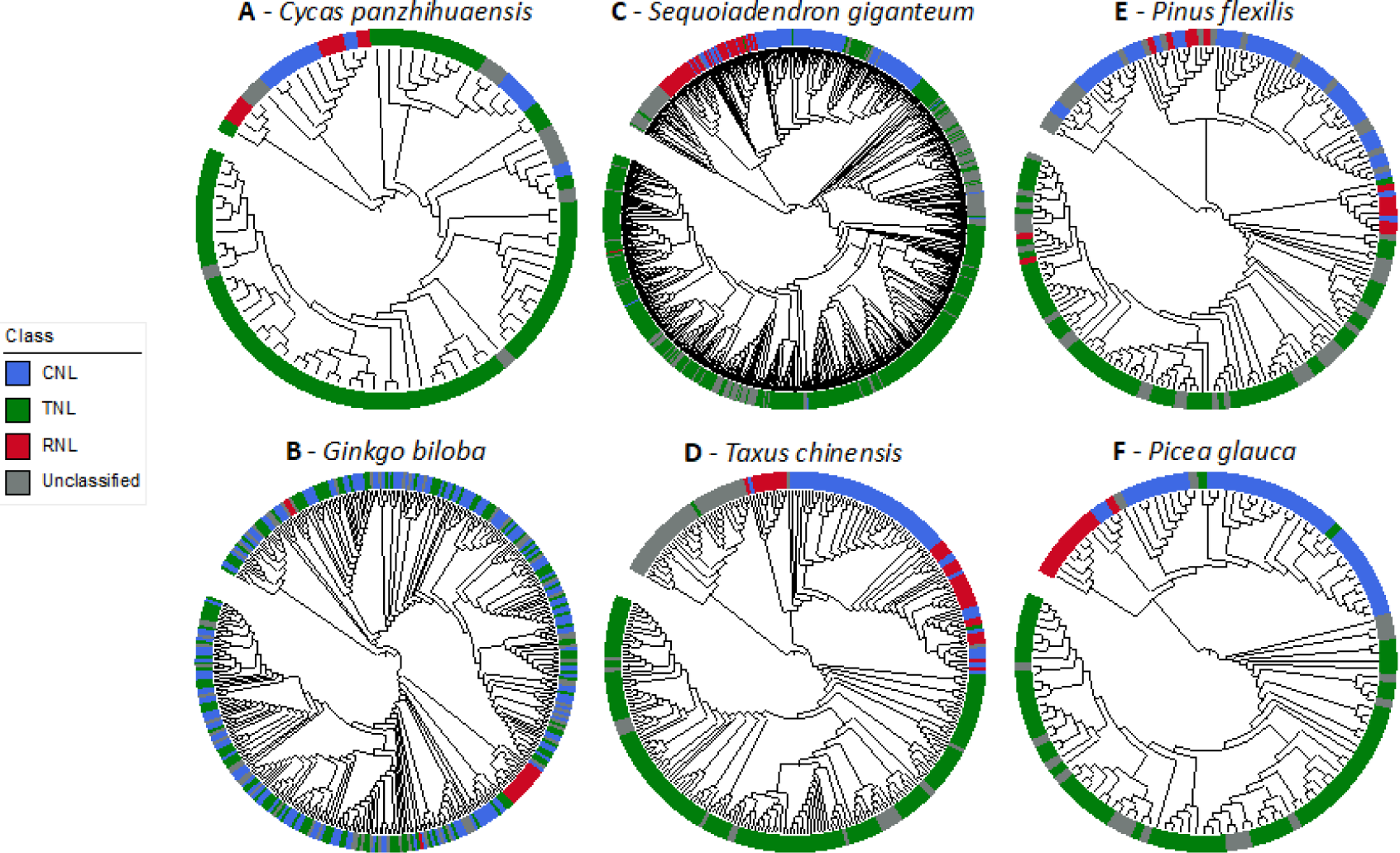
Intragenomic phylogenetic relationships of NLR genes based on the conserved central NB-ARC domain, calculated with maximum likelihood algorithms using IQTree v1.6.12 (Nguyen et al. 2015) (Materials & methods section 3) and annotated using the iTOL web server (Letunic and Bork 2021). Colour strips around the circular trees indicate NLR class as determined with NLR Annotator (Steuernagel et al. 2020) and manual BLASTs (Materials & methods section 2). Main gymnosperm lineages represented by Cycadales (A), Ginkgoales (B), Cuppressaceae (C), Taxaceae (D), and Pinaceae: *Pinus* (E), and *Picea* (F).

The densest clusters in conifers are mainly composed of TNLs (Fig. 2B-E), which correlates with their intrachromosomal diversification (Fig. 4G). Most TNLs that occur on the same chromosome are also phylogenetically correlated. In the Pinaceae this phylogenetically correlated diversification even occurred on the same syntenic linkage group (Supplementary Materials, Figure S1-S4).

## Discussion

The study of NLR immune receptor evolution and genomic architecture in conifers is motivated in part by the increased threats from diverse biotic aggressors. The 615 species of extant conifers span all continents except Antarctica and are classified in eight families with the largest being Pinaceae, Cupressaceae, and Podocarpaceae (Farjon and Filer 2013). Forest trees, including conifers, have several co-evolved biotic aggressors ranging from rust diseases to herbivorous insects which may damage or even kill trees. However, more severe attacks, range expansions or species introductions, and infection of previously unknown hosts have become increasingly prevalent and linked to climate change (Teshome, Zharare, and Naidoo 2020). Major herbivorous insects are expanding their range and intensifying damage levels in conjunction with climate change, such as mountain pine beetles (e.g. Mitton and Ferrenberg 2012). New diseases or outbreaks in several conifers are being linked to Oomycetes, namely species of *Phytophthora* de Barys (e.g., Brasier and Webber 2010, Brar et al. 2018). There is growing evidence of the involvement of TNLs in resistance particularly to rust diseases affecting pines (Quesada et al. 2014; Amerson et al. 2015; Ence et al. 2022), but much less is known when it comes to insects and newly emerging biotic threats. Understanding the genomic architecture of NLR genes will therefore be critical for informing breeding programs and other conservation practices seeking to mitigate the effects of climate change.

### Ubiquitous genomic clustering of NLR genes in conifers

Through comparative genome-wide analysis of NLR immune receptor genes in conifers, we elucidated a conserved non-random chromosomal distribution of NLR genes. In all conifer genomes analysed here we found one chromosome containing a disproportionately large number of NLR genes (Figure 2). Without exception, these chromosomes contained dense clusters of NLR genes (Figure 3), comprising on average a quarter of the total number of NLR genes in the genome (Figure 2B-E). These clusters predominantly contain TNL genes, which share an ancestral origin (Figure 4 & 5). High-density gene regions are a hallmark of conifer genomes (Pavy et al. 2017), hinting at tandem duplications, as detected frequently for NLR genes (Pavy et al. 2017) and for other conifer gene families (e.g. Guillet-Claude et al. 2004). We therefore conclude that the unbalanced genomic clustering pattern of NLR genes found previously in *Pinus flexilis* (J.-J. Liu et al. 2019) occurs in a taxonomically broad range of other conifer species. The absence of this pattern in non-coniferous gymnosperms (Figure 3 & Figure 4A-B) indicates that NLR gene clusters arose in the ancestor of conifers. Diversification within these large groups of sequences appears to be variable and largely lineage-specific (such as in the Pinaceae). Partial functional redundancy is likely, in response to similar environmental cues and selection pressures among conifer taxa, thus contributing to the high abundance of NLR genes in conifers. These ancestral clusters may have enabled lineage-specific tandem duplications leading to the high (and variable) abundance of NLR genes observed in conifers (Van Ghelder et al. 2019; this study), as compared to non-coniferous plants such as cycads (Table 1, Figure 4A) and many angiosperms. A high diversity of NLR genes may lead to a high versatility in drought and disease resistance for these woody perennials characterised by delayed sexual maturity for reproduction. These genes are likely to contribute to the ecological dominance of conifers in many boreal and temperate forests (Bonello et al. 2006), including in highly inhospitable habitats (Laberge et al. 2000).

Genomic clusters of NLR genes have previously been found in a variety of angiosperm lineages (Wersch and Li 2019), such as *Arabidopsis* Heinh. (Brassicaceae) (Meyers et al. 2003), lettuce (Asteraceae) (Christopoulou et al. 2015), peach (Rosaceae) (Verde et al. 2013), potatoes (Solanaceae) (Seo et al. 2016), and wheat (Poaceae) (Smith et al. 2007). In contrast to the studied conifers, NLR clustering is frequent but not as ubiquitous in angiosperms. What further sets conifers apart from angiosperms in regards to NLRs, is the ubiquitous presence and abundance of all three NLR subfamilies (CNLs, RNLs and TNLs) (Van Ghelder et al. 2019, Table 1 of this study). RNL abundance is rare in angiosperms and TNLs are absent in monocots (Van Ghelder et al. 2019). We found RNLs comprised 4-14% of the total NLR diversity in all gymnosperms, indicating that RNL abundance is an ancestral trait to the gymnosperms. RNLs were consistently divided over two phylogenetic clades, one of which comprised mainly CNLs (Figure 4). This is consistent with an evolutionary divergence between TNLs and CNL/RNLs predating that between CNLs and RNLs (Shao et al. 2019). Interestingly, CNLs and TNLs shared ancestral origins in *Ginkgo biloba* (Figure 4B). This potentially indicates frequent domain swapping between NLR genes, highlighting the dynamic nature of these resistance genes.

### Evolution of genomic architecture in conifer giga-genomes

Conifers have very large genomes (18-34 Gb) and harbour large amounts of repetitive DNA sequence (Mackay et al. 2012; Nystedt et al. 2013; Birol et al. 2013; Zimin et al. 2014; De La Torre et al. 2014). Genome evolution is considered to be less dynamic in conifers and other gymnosperms compared to flowering plants (Leitch and Leitch. 2012), which may suggest lower rates of gene diversification. On the other hand, conifers have suites of rapidly evolving genes (Gagalova et al. 2022) and highly diversified gene families or sub-families (e.g., Bedon et al. 2010; Stival Sena et al. 2018; Van Ghelder et al. 2019) both related to stimuli and stress response. Several comparative studies in conifers have shown high levels of intergeneric macro-synteny and macro-collinearity among *Pinaceae* taxa (Pavy et al. 2012) (Pelgas et al. 2006; Ritland et al. 2011; Westbrook et al. 2015) and clear chromosomal rearrangements when comparing Pinaceae and Cupressaceae (Moriguchi et al. 2012, de Miguel et al. 2015). These observations are consistent with a small number of whole genome duplications early in conifer evolution (Li et al. 2015). In contrast, our study has focused on the genomic architecture of a targeted gene family, showing conserved localised clustering and shedding insights into the evolutionary trajectory of NLR genes. Our investigation was possible due to two types of relatively recent genomic resources: 1) highly contiguous genome assemblies, e.g., *Sequoiadendron giganteum* (Lindl.) J.Buchholz (Scott et al. 2020), *Taxus chinensis* (Pilg.) Rehder (Xiong et al. 2021), among others, which are developed using proximity ligation (e.g., HiC (Belton et al. 2012)) and long-read sequencing (e.g., PacBio SMRT); 2) high density genetic maps which are available for *Pinus* L. *spp.* (e.g., J.-J. Liu et al. 2019), and *Picea* A.Dietr. *spp.*(Bernhardsson et al. 2019; Gagalova et al. 2022; Tumas et al. 2023).

A specific feature of conifer genomes probably enabled the accumulation of NLR genes and facilitated the formation of very large gene clusters. Conifers are inefficient at removing extra copies of DNA sequence through proof-editing, hence their propensity to accumulate gigabases of repetitive sequences such as the type I transposable elements (e.g. copia and gypsy sequences) (Nystedt et al. 2013; Zimin et al. 2014). Conifers also retain a high proportion of pseudogenes (Warren et al. 2015) and single copy sequences that are similar to protein coding genes (Pellicer et al. 2018). Therefore, we could expect a significant proportion of genomic NLR sequences in conifers to represent pseudogenes. However, RNA sequencing has identified between 271 and 725 NLR genes expressed across a suite of Pinaceae and Cupressaceae species (Van Ghelder et al. 2019; Liu et al., 2021; Ence et al., 2022). Expression of selected NLRs was shown to be responsive to infection by *Phytophthora ramorum* in *Larix spp*. (Dun et al. 2022) or drought in *Picea glauca* (Van Ghelder et al. 2019) and to be variable across different seed families in *Pinus flexilis* (J.-J. Liu et al. 2021). The genomic sequences identified here may prompt further work to determine which of these are expressed and under what circumstances. Studies of functional divergence among sequences and of evolutionary rates, aiming at identifying footprints of natural selection (e.g., Guillet-Claude et al. 2004) should also be considered (e.g., Chia and Carella 2023).

### Opportunities for genomic breeding using NLR genes for pest, disease and drought resistance

Resistance to viruses, bacteria, oomycetes, fungi and some insects has been linked to NLR genes in a range of flowering plants (Kourelis and van der Hoorn 2018) but our understanding of their contribution to resistance in conifers is very rudimentary. Dissection of the genetic resistance to fusiform rusts (*Cronartium quercuum* (Berk.) Miyabe ex Shirai f.sp. *fusiforme*) in *Pinus taeda* L. breeding populations has provided evidence for genetic resistance (Wilcox et al. 1996) and implicated NLR-encoding genes as likely candidates (Quesada et al. 2014; Ence et al. 2022). In *P. taeda,* nine different fusiform rust resistance loci were identified across three linkage groups (Amerson et al. 2015). These are hypothesised to contain NLR sequences, which were found to vary in numbers of genomic sequences when comparing populations from different geographic areas (Ence et al. 2022). Similarly, in *Pinus flexilis*, fine genetic dissection, evolutionary analysis, and expression profiling have identified two NLR genes as candidates for resistance linked to the Cr4 locus (J.-J. Liu et al. 2021) among 155 NLR genes mapped across 12 linkage groups to date. Interestingly, the major clusters found in *P. flexilis* were on linkage groups distinct to the Cr4 locus. Taken together, these results indicate that resistance-associated NLR genes may be distributed across the genome. To our knowledge, resistance phenotypes have been identified in only a few studies in conifers and these have involved fungal rusts infecting pines, although expression profiling indicated responsiveness to *Phytophthora ramorum* (Dun et al. 2022) and drought in other species (Van Ghelder et al. 2019).

NLR-dense genomic regions could act as potential reservoirs for NLR diversity. The NLR clusters found in gymnosperm genomes probably arose by tandem duplications of TNL genes, as indicated by their consistent composition in conifers (Figure 2B-D) and the close phylogenetic relationships of the TNLs on the same chromosome (Figure 5). Although tandem duplication inevitably leads to identical gene copies initially (paralogs), it can eventually lead to gene diversification through differential mutation trajectories and domain swapping (Ostermeier and Benkovic 2001). There are documented examples of such neofunctionalisation for resistance genes in different plant lineages (Kong and Ranganathan 2008; Wei et al. 2023) and for genes encoding transcription factors in the Pinaceae (Guillet-Claude et al. 2004). Given the long evolutionary history of conifer lineages (Leslie et al. 2018) and their considerable NLR diversity, a high degree of non-canonical NLR genes is expected, especially in and around these dense NLR gene clusters. Considering whole-genome datasets, about one in four NLR genes is unclassified in conifers (Table 1), meaning that a distinctive N-terminal domain is missing or ambiguous (e.g., fusion of TIR and CC domains). Closer inspection of NLR-dense regions could therefore reveal interesting non-canonical domains in conifer NLR genes. Together with the overall diversity of canonical NLR genes in these clusters, this could further emphasise their potential in genomics-assisted breeding to improve disease and drought resistance.

**Figure 5:**
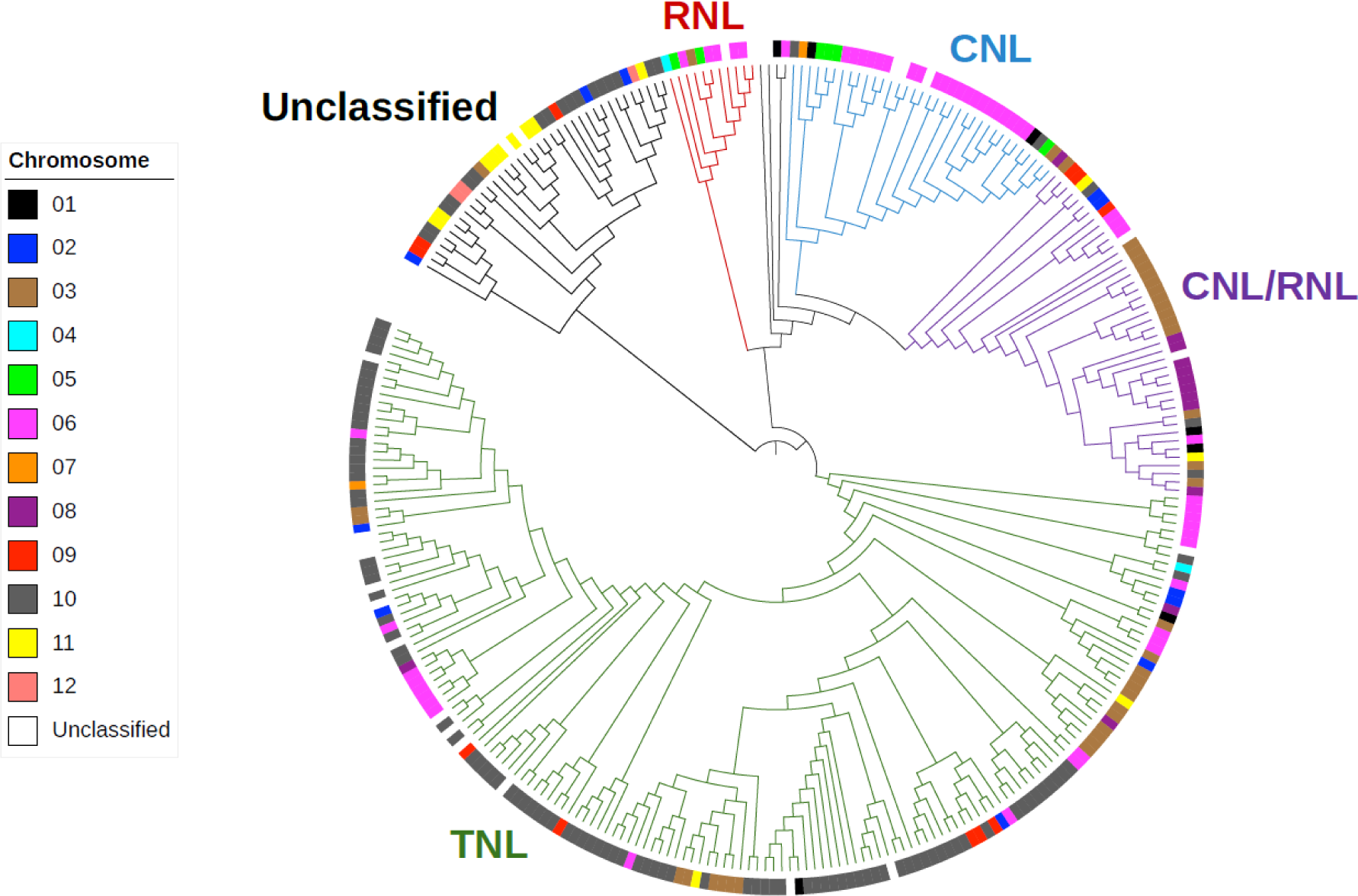
Chromosomal structuring in phylogenetic relationships within a conifer species (*Taxus chinensis*) displayed in an intragenomic framework. Branch colours correspond to the NLR gene subfamily, as indicated in the same colours next to the phylogeny. Coloured squares at the tips of branches represent the chromosome on which the respective NLR genes are located. NLR genes found on scaffolds that were not assembled into chromosomes are indicated with empty colour squares (“Unclassified”).

The NLR gene family is among the most studied in plants due to its agronomic importance (Kourelis and van der Hoorn, 2018) with many linked breeding and genetic engineering applications proposed to improve resistance in crops (van Wersch et al. 2020) and to a lesser extent in forest trees (Ence et al. 2022). We have shown how improved genome sequences along with transcriptome data may enhance our understanding of NLR genomic architecture in this understudied group. In order to develop genetic resources that will help to respond to emerging threats in conifers, three other components are needed: 1) populations of phenotypically diverse individuals in which to study resistance traits (Ence et al. 2022; Liu et al., 2019); 2) efficient and accurate assessment of the susceptibility and resistance phenotypes to relevant pests and diseases, which is difficult to accomplish and is therefore often either lacking or sub-optimal in conifers; 3) fast and accurate genome scanning methods that are suitable for differentiating among genes and alleles within and among populations. The large size and the variability of the NLR gene family adds to the challenge of linking genes to resistance phenotypes; however, knowledge of the position of clusters of these immune receptor genes paves the way to more focused investigations in conjunction with genome selection and other genome-wide analyses.

## Materials & methods

### 1 Genomic distribution of NLR genes in Pinaceae and other conifer families

Following the evidence for dense genomic clusters of NLR immune receptor genes in *Pinus flexilis* (J.-J. Liu et al. 2019), we tested whether this is a Pinaceae family-wide phenomenon by examining publicly available genomic resources in other members of the family. To characterise the distribution of NLR genes in the speciose *Picea* genus, we deployed a high-density linkage map for *P. abies* (Bernhardsson et al. 2019) and high-quality genome assemblies of *P. glauca* and *P. sitchensis* (Gagalova et al. 2022). Although these assemblies do not have chromosome-level contiguity, they are suitably scaffolded into linkage groups corresponding to an updated version of the original *P. glauca* high-density linkage map (Pavy et al. 2017).

NLR genes were identified using the NLR Annotator pipeline v2.1 (Steuernagel et al. 2020) and linkage map positions were recorded to map NLR gene distribution on the twelve different linkage groups. We utilised the recent linkage map comparison work by Tumas et al. (2023) to determine the syntenic linkage groups. NLR distribution data for *P. flexilis* were taken from the original high-density linkage map publication (J.-J. Liu et al. 2019).

To compare distribution patterns across main lineages of conifers and gymnosperms, we applied the same pipeline to recently published chromosome-level genome assemblies: *Sequoiadendron giganteum* (Cupressaceae) (Scott et al. 2020), *Taxus chinensis* (Taxaceae) (Xiong et al. 2021), *Ginkgo biloba* (Ginkgoales) (H. Liu et al. 2021), *Pinus tabuliformis* (Niu et al. 2022), and *Cycas panzihuaensis* L.Zhou & S.Y.Yang (Cycadales) (Y. Liu et al. 2022). Nucleotide positions of NLR genes were recorded for each chromosome.

To visualise the distribution of NLR genes, histograms were produced in R (R Core Development Team 2010) with ggplot2 v3.4.2 (Wickham 2016) where genes were plotted along the length of chromosomes (linkage groups for linkage maps) using the starting position in Mbp (cM for linkage maps). The bin width was chosen to correspond roughly to 1% of the largest chromosome (or linkage group), i.e.: 13 Mbp for *C. panzihuaensis,* 12 Mbp for *G. biloba*, 10 Mbp for *S. giganteum* and *T. chinensis*, 4 Mbp for *Picea glauca*, 2 Mbp for *P. sitchensis* (Bong.) Carrière, 4 cM for *P. abies* (L.) H.Karst., and 2 cM for *Pinus flexilis*. We calculated the expected number of NLR genes for each chromosome (or linkage group) to quantify abnormal distribution patterns:

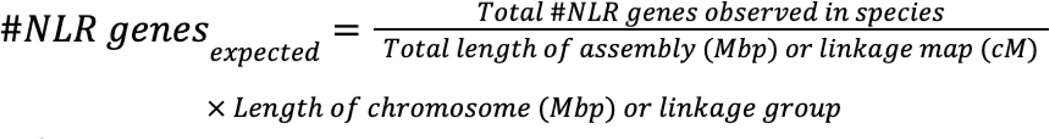

The anomalies were visualised by plotting the observed values against the expected values in a scatter plot. To determine whether these anomalies were statistically significant we performed two-character Fisher’s exact tests in R (R Core Team 2021) and computed the p-values.

### 2. NLR classification

The NLR Annotator pipeline describes the motifs discovered in each NLR based on a curated list of NLR motifs (Jupe et al. 2012). We utilised a custom python script (available on https://github.com/hung-th/NLRmeta), adapted from open-source code by Philipp Bayer (https://gist.github.com/philippbayer/0052f5ad56121cd2252a1c5b90154ed1) and based on the motif table in Jupe et al. (2012), to extract the motif output from NLR Annotator and convert it into CNL or TNL subfamily classification. The third subfamily, RNL, is characterised by the variable N-terminal RPW8 domain but is not annotated by NLR Annotator. In a previous study on conifer NLR genes, Van Ghelder et al. (2019) discovered two RNL-characteristic motifs located in the RNBS-D domain. We searched for these motifs in the generated NLR datasets by deploying the MAST software from the MEME suite (Bailey et al. 2015) and classified NLR genes containing these as RNLs. A further search for RPW8 domains was performed with tblastn (eV <0.05) using the conifer RPW8 sequences characterised by Van Ghelder et al. (2019) against the genome assemblies and linkage map loci. We only retained RPW8 hits with ≥1/3 amino acid identity in the reference sequences. NLRs with an RPW8 domain fused to the N-terminal side were thereby classified as RNLs regardless of their RNBS-D motif composition. Separate RPW8 sequences (e.g., not fused to an NLR gene) were also characterised as RNL genes. A fourth category of “unclassified” NLRs encompasses NLR genes that could not be classified into one of the three subfamilies due to a lack of characteristic domains and motifs. NLR genes containing motifs characteristic of CNL as well as TNL were also labelled as “unclassified”. For *P. flexilis*, we utilised the annotation information from the original linkage map publication (J.-J. Liu et al. 2019) to divide NLRs into classes. RNLs in *P. flexilis* were classified in the same way as for the other species. NLR class information was used to further annotate the intrachromosomal NLR distribution histograms (Materials & methods section 1) to visualise patterns of class distribution.

NLR Annotator further determines whether detected NLR genes are complete, partial (missing domains) or pseudogenes (unexpected stop codon in sequence), which we recorded for each discovered gene, except for the separate RPW8 sequences.

### 3. NLR phylogenies

To determine the evolutionary history of discovered NLR genes, maximum likelihood phylogenies were generated from alignments of the central (conserved) NB-ARC domain. For *Pinus flexilis*, NB-ARC sequences were identified through a BlastP search (eV <0.05) of the reference NB-ARC sequence used by NLR Annotator against the translated NLR sequences. All hits with ≥60 amino acid residues were extracted and aligned using MAFFT v7 (Katoh and Standley 2013) using default settings. For the other species, we utilised the ‘-a’ flag in NLR Annotator to obtain NB-ARC domains of all complete NLR genes. Maximum likelihood phylogenetics was performed with IQTree v1.6.12 (Nguyen et al. 2015) using the ‘GTR20’ model for protein evolution and 1000 ultra-fast bootstrap replicates to calculate node support values. Phylogenies were visualised in the online Interactive Tree of Life (iTOL) tool (Letunic and Bork 2021) and rerooted at the node separating RNL/CNL and TNL clades. NLR class and chromosome were mapped onto the topologies using the iTOL annotation editor.

## Acknowledgements

We are grateful for feedback to this study and its results from the Sitka Spruced (https://sitkaspruced.web.ox.ac.uk/) and Spruce-Up communities (https://spruce-up.ca/en/). As part of the Sitka Spruced project (BB/P020488/1), this research was funded by BBSRC and a group of forest and wood processing industries.

## Author contributions

YW & JJM designed the study. YW conducted the bioinformatic analyses with input from HT, CVG & THH. YW & JJM wrote the manuscript with input from all authors.

## Data availability

Genomic mapping data of NLR gene distributions generated for this study is available in the Figshare digital repository under the following DOI: 10.6084/m9.figshare.24412579

## Supplementary Materials

A detailed description of the phylogenetic analysis of NLR genes in Pinaceae is available in the Supplementary Materials. The following figures are available in this document:

Fig. S1: NLR phylogeny for Pinus flexilis.

Fig. S2: NLR phylogeny for Picea abies.

Fig. S3: NLR phylogeny for Picea glauca.

Fig. S4: NLR phylogeny for Picea sitchensis.

