## Supporting Materials for "Conifers concentrate large numbers of NLR immune receptor genes on one chromosome"

**Description:** Details of the phylogenetic analysis of NLR genes in Pinaceae are described here. NLR genes were mapped onto the genetic linkage maps and associated genome assemblies available in *Pinus flexilis* (J.-J.Liu et al. 2019), *Picea abies* (Bernhardsson et al. 2019, *G3*), *Picea glauca* (Pavy et al. 2017, *The Plant Journal*; Gagalova et al. 2022, *The Plant Journal*) and *Picea sitchensis* (Gagalova et al. 2022, *The Plant Journal*). Synteny between linkage groups was determined using the recently produced genetic linkage map for *Picea sitchensis* (details in Tumas et al. 2023, *BioRXiv*, https://doi.org/10.1101/2023.08.21.554184). NLRs are analysed in a maximum likelihood phylogenetic context based on alignments of the conserved central NB-ARC domain (details in Methods section 3, main text). Tips of topologies were annotated for NLR class (colour strip) and linkage group (symbol). Linkage groups were numbered according to the linkage map for *Pinus flexilis*, based on synteny. In all species TNL diversification was prolific on linkage group #02.

**Fig. S1**: NLR phylogeny for *Pinus flexilis*.
**Fig. S2**: NLR phylogeny for *Picea abies*.
**Fig. S3**: NLR phylogeny for *Picea glauca*.
**Fig. S4**: NLR phylogeny for *Picea sitchensis*.

**Fig. S1**

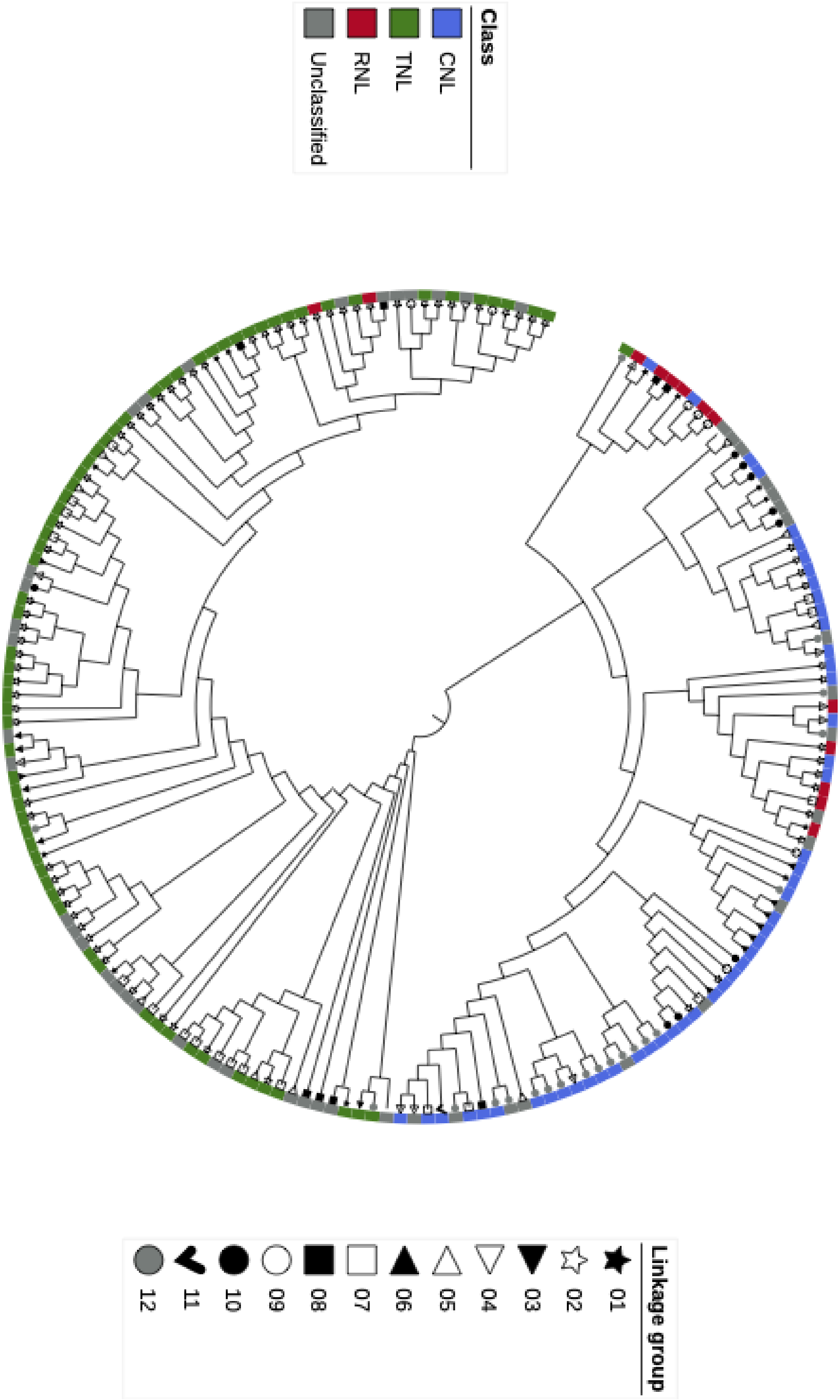
Class
CNL
TNL
RNL
Unclassified
Linkage group
01
02
03
04
05
06
07
08
09
10
11
12

**Fig. S2**

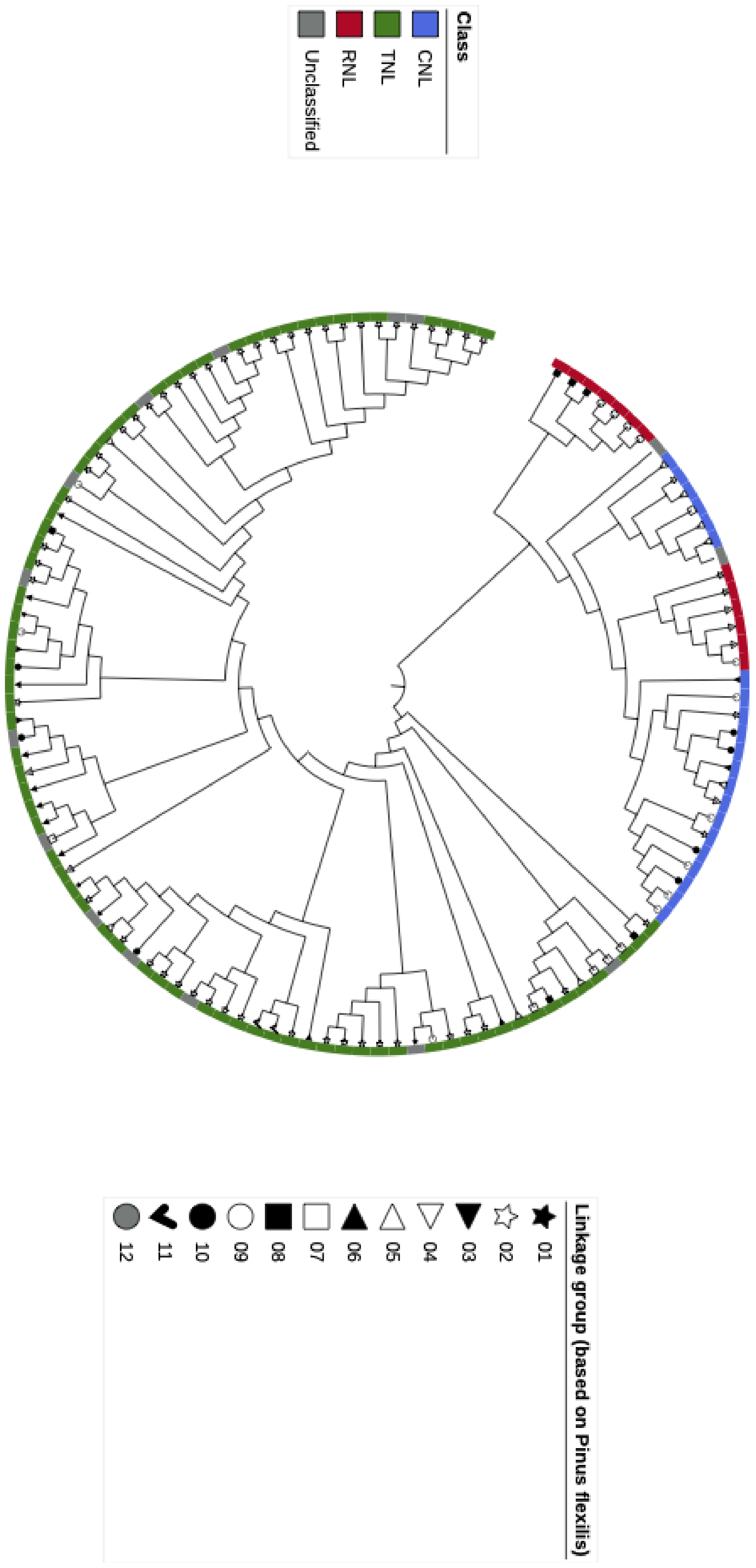

Class
CNL
TNL
RNL
Unclassified
Linkage group (based on Pinus flexilis)
01
02
03
04
05
06
07
08
09
10
11
12

**Fig. S3**

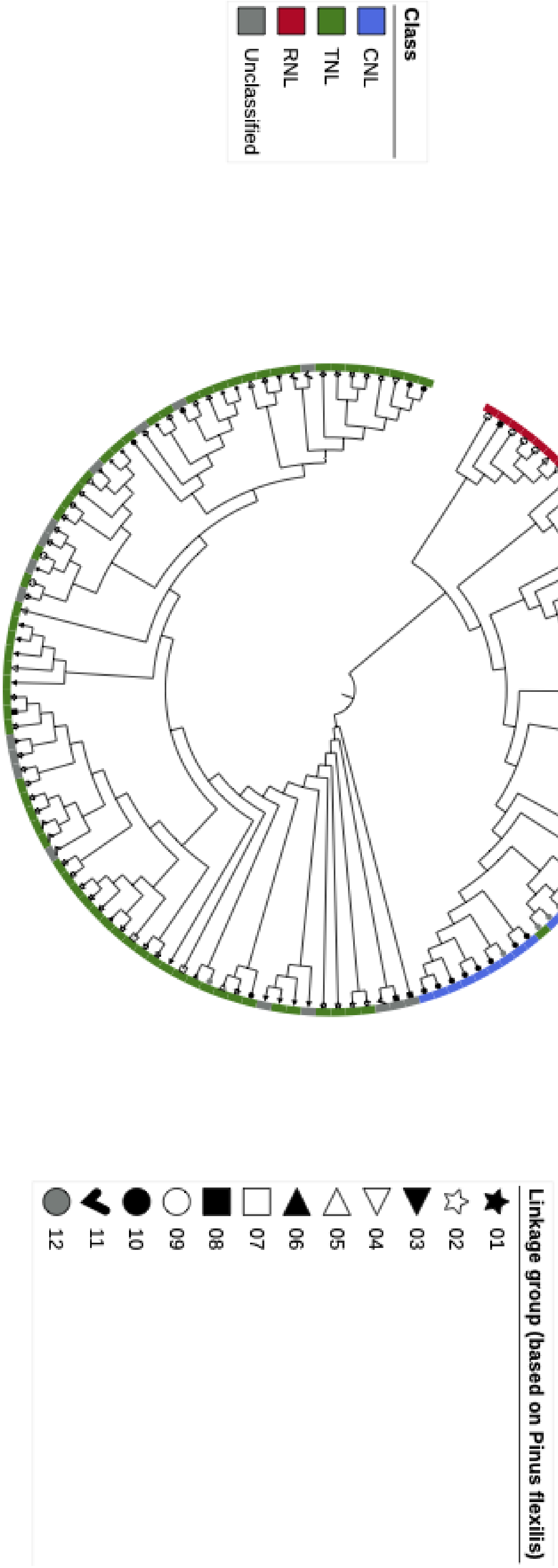

Class
CNL
TNL
RNL
Unclassified
Linkage group (based on Pinus flexilis)
01
02
03
04
05
06
07
08
09
10
11
12

**Fig. S4**

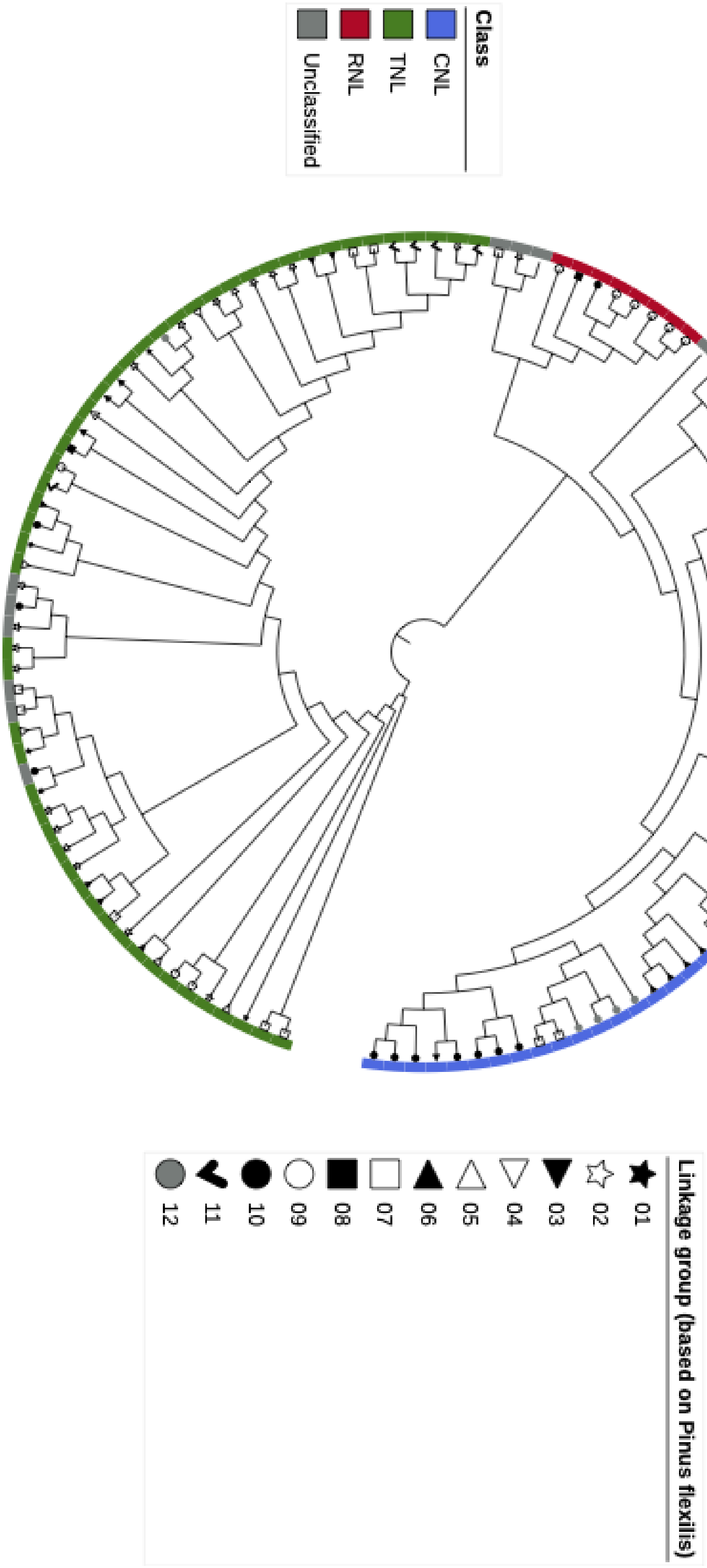
Class
CNL
TNL
RNL
Unclassified
Linkage group (based on Pinus flexilis)
01
02
03
04
05
06
07
08
09
10
11
12
